# Harnessing the power of metabarcoding in the ecological interpretation of plant-pollinator DNA data: strategies and consequences of reads filtering

**DOI:** 10.1101/2021.06.14.448412

**Authors:** Nicola Tommasi, Andrea Ferrari, Massimo Labra, Andrea Galimberti, Paolo Biella

**Author notes:** Correspondence P.za Della Scienza, 2 20126-I, Milan, Italy.

## Abstract

DNA metabarcoding approaches to analyse complex mixtures of pollen has become the standard in pollination biology, especially in the light of recent threats affecting pollination. In spite of the increasing adoption of High Throughput Sequencing (HTS) approaches, these studies generate huge numbers of raw reads, some of which might be associated to false positives or infrequently recorded species with potentially little biological information. If these reads are not discarded (i.e. pruned), they can lead to changes in the ecological findings and lead to biased conclusions. In this study we reviewed 42 papers in the recent pollen DNA metabarcoding literature and focused on the type of pruning applied. We also tested whether the different types of those cut off threshold may leave a mark on the DNA metabarcoding data. To do so, we compared for the first time community composition, species richness and networks of species interactions (i.e. Connectace, Modularity, Connectivity and Shannon entropy) associated with the most relevant ways of treating HTS outputs: no cut (no reads filtering), or cutting levels obtained as proportional 1% of sample total reads, or as fixed amount of 100 reads, or from ROC (Receiver operator characteristic). Results clearly indicated that pruning type shapes species composition and that to apply or not a threshold dramatically impacts ecological indices, potentially increasing the risk of misinterpreting DNA metabarcoding data under an ecological point of view. Given the high methodological heterogeneity from the revised literature, we discuss in what conditions filtering types may be more appropriate, and also recommend to biologically justify the pruning threshold when analysing DNA metabarcoding raw reads, and to develop shared approaches to make future studies more comparable.

## 1- Introduction

The study of plant-pollinator interactions is pivotal to address both theoretical and applicative issues at the global scale, with important implications in evolutionary studies, conservation biology, agrifood security and to provide reliable policies of land-use management and mitigation of anthropogenic stressors (Mitchell et al., 2009; Schweiger et al., 2010; Burke et al., 2011; Burke et al., 2017).

Traditionally, studies of plant-pollinator interactions have been carried out through direct field observations of insects foraging activity while visiting flowers (CaraDonna & Waser, 2020; De Manincor et al., 2020). However, another valuable approach to unveil information on plant-pollinator interactions is based on the identification of the pollen grains that the pollinator insects carry on their body (Bosch et al., 2009; Cullen et al., 2021). While visiting flowers, pollinators get in touch with the flower’s anthers or actively collect and accumulate the pollen in specialized structures such as the *scopa* or the *corbiculae*. The characterization of the transported pollen allows shedding light on the foraging ‘history’ of insects prior to a sampling event. In this way, it is possible to retrieve complete behavioral and ecological information on flower resource exploitation, and to properly address ecological research questions. The palynology approach has traditionally been used to identify pollen samples, and it requires high expertise with light-microscopy-based species assessment and is time-consuming (Bell et al., 2016 (a), Bell et al., 2016 (b)). In addition, reaching a detailed taxonomic resolution through morphological criteria could be limited by the lack of diagnostic characters among congeneric species (Khansari et al., 2012).

In the last decade, these difficulties have progressively been addressed due to the increasingly accessible DNA-based identification technologies that significantly reduced the time required for pollen identification (Galimberti et al. 2014, Bruni et al., 2015). Recent developments in DNA sequencing technologies, such as the increasing adoption of High-Throughput Sequencing (HTS) facilities, made it possible to analyse the taxonomic composition of complex DNA matrices, including pollen (Liu et al., 2012), using standard DNA barcode regions in a so-called DNA metabarcoding approach (Taberlet et al., 2012). In the field of pollen-based studies, the use of DNA metabarcoding soon become a standard approach and to date, it has been employed not only in the characterization of the pollen retrieved from insects bodies (see e.g. Biella et al, 2019), but also in the analysis of other kinds of matrices, such as the pollen stored in cavity nests (McFrederick et al., 2016), honey (Richardson et al., 2015 (a)), sediments (Niemeyer et al., 2017; Alsos et al., 2018) and was also employed in the fields of honey authentication (Bruni et al., 2015; Prosser & Hebert, 2017) and forensics sciences (Ezegbogu, 2021, Bell et al 2016 (b)). In the context of plant-pollinator interactions, the data retrieved from pollen DNA metabarcoding could potentially shed light on how pollinators exploit flower resources and consequently to evaluate the complexity and resilience of the interaction networks in a given habitat. This methodological revolution not only improved ecological knowledge, but also offered new insights into the development of effective conservation and restoration actions (Bell et al., 2016 (a)). Given the astounding number of sequences (hereafter “reads”) obtained through HTS techniques, (Churko et al., 2013; Bell et al., 2019) it is necessary to process the data through a proper bioinformatic pipeline. This is a critical phase of the dry lab activities and usually consists in (i) the assembly of paired-end reads resulting from bidirectional sequencing of the DNA templates, (ii) the analysis of the variation among sequences and the clustering of molecular features (e.g., Operational Taxonomic Units OTUs *sensu* Blaxter et al., 2005 or exact sequence variants ESVs *sensu* Callahan et al., 2017), and finally (iii) the removal of chimeras, artifacts and spurious sequences (Alberdi et al., 2018). However, this process does not solve the biases that could be introduced at different stages of the DNA metabarcoding workflow and that could alter the species detection. These include for example the choice of primer that could preferentially amplify certain taxa during PCR (as highlighted in Piñol et al., 2019), to the choice of the clustering method for calculating the molecular features (Clare et al., 2016). Species characterization could also be altered by the application of a cut-off threshold, usually applied to remove those reads resulting from rare occurrence and/or potential contaminants, and this is likely the most crucial step where severe biases in terms of species composition of the investigated biological matrix can be introduced (Ficetola et al., 2016; Alberdi et al., 2018). HTS technologies have the potential to magnify the sequencing errors and overestimate the presence of rare species, thus the application of a threshold that balances the detection of rare reads and removing artifacts is of particular importance (Alberdi et al., 2018). These artifacts include for example the false positives that are clusters of molecular features (i.e., OTUs and ESVs) generated as a consequence of inaccuracies during field sampling operations (e.g cross-contamination among samples), laboratory processing (e.g., contamination of DNA extraction or amplification reagents), or bioinformatics analysis (e.g., misidentification or maintenance of chimeric sequences) (Ficetola et al., 2016; Bell et al., 2019). Therefore, the extreme sensitivity of DNA metabarcoding approaches makes it crucial to filter out false positives and rare occurrences during the post-sequencing bioinformatics processing. A typical problem in studies that use DNA metabarcoding related to pollination biology could derive for example from the identification of plant pollinator interactions that actually have never occurred in the field, or that are accidental or extremely rare. The characterization of plant taxa from pollen samples could also depend on rare pollen occurrences (i.e a single pollen grain) whose presence led to ecological interpretation consequences, especially when reads count are converted and used as presence absence data. These, in turn, could lead to the overestimation of the generalist attitudes of the investigated insects and therefore to misleading ecological interpretations.

The application of an appropriate “cut-off threshold” to “prune” the DNA metabarcoding data from the signal of possible false positives and rarest occurrence is therefore a critical step of the bioinformatics pipeline. Although some studies did not apply any cut-off threshold, different types of pruning have been used so far in recent literature. This highlights the absence of agreement on whether and how to prune a DNA metabarcoding output. In practice, some studies are based on fixed cut-off thresholds, such as a defined number of reads used as reference level for accepting a molecular feature in a sample (Pornon et al., 2019). Other studies employed proportional cut-off thresholds, where molecular features are discarded if represented by less than a certain percentage of the total reads produced for a sample (Wilson et al., 2021). Alternatively, statistical approaches have been used for estimating a variable threshold based on Receiver operator characteristic (ROC) curves, thus depending on the distribution of reads among molecular features within a sample (Biella et al., 2019). However, to date, no studies have investigated the effect of different cut-off thresholds on molecular datasets, specifically from studies related to mixed pollen samples (or honey) and plant-pollinator interactions.

In this study, we investigated the criteria adopted for pruning the false positives and rarest occurrence in published pollen DNA metabarcoding studies, first by summarising the strategies on the application of the cut-off threshold for false positives removal in the recent scientific literature. Moreover, we aimed at evaluating the direct ecological effects of the most commonly applied false positive removal methods on publicly available pollen/honey DNA metabarcoding datasets. To do this, we measured how different cut-off thresholds impacted i) species composition and species richness in the samples, and iii) the interactions among plants and pollinators described by network indexes calculated at both at the community and the individual level. This approach allowed us to evaluate how the different pruning strategies could alter the identification of species and thus the ecological interpretation of results.

## 2- Methods

### 2.1 - CUT-OFF THRESHOLD APPLICATION IN POLLEN DNA METABARCODING: LITERATURE OVERVIEW

To summarise the typology of cut-off threshold already applied in the scientific literature a bibliographical research was conducted in Scopus using the following keywords: “DNA” + “metabarcoding” + “pollen”. Within the results, we selected only peer-reviewed original published articles that dealt with pollination, pollinator diet (pollen and honey), and plant pollinator interactions with a DNA metabarcoding approach. We excluded reviews, news, views, opinions, and perspectives papers, keeping only research articles based on original data. Papers on airborne pollen or other pollen matrices have been excluded too because unrelated to pollinators. We selected studies spanning between 2012, when the term DNA metabarcoding was proposed for the first time (Taberlet et al., 2012) and 2021, with the last update on the 9th of May 2021. The retrieved articles were used for the creation of a review table to summarise the following information: (i) the type of sample from which the DNA was extracted, (ii) the studied organism, (iii) the details of the post-sequencing cut-off threshold applied in the analysis pipeline, and (iv) the DNA barcoding markers used to achieve the amplification reaction.

### 2.2 EVALUATING THE CONSEQUENCES OF CUT-OFF THRESHOLDS APPLICATION

To evaluate how the application of different cut-off thresholds could lead to changes in the results obtained through DNA metabarcoding of mixed pollen samples, publicly available DNA metabarcoding datasets (obtained by ITS2 DNA barcode marker sequencing, the most recurrent marker in pollen DNA metabarcoding studies) were retrieved from the previously mentioned literature search. Only datasets containing a non-filtered number of reads were kept for our analysis (see Table 1 and Results). In detail, we retrieved published non filtered dataset (hereafter named as “no cut”, equivalent to a 0-reads threshold), and we derived several subsequent “pruned” versions by applying each of three (independently) different approaches to calculate the reads cut-off thresholds. The chosen pruning types were based on utilization frequency in the literature, or if based on promising approaches of biological importance (i.e., the ROC approach). Specifically, the first method is proportional and discards molecular features represented in a sample with a number of reads lower than 1% of the total sample reads count (hereafter “proportional 1%”) as used in Danner et al., 2017. The second one estimates a cutting threshold accounting for the distribution of reads among molecular features, thus providing a customized proportion for each sample through the statistical ROC curve approach, as indicated in Biella et al (2019) (hereafter “statistical ROC”). This strategy is commonly applied in several disciplines and was specifically proposed for false positive detection (Metz, 1978), and it bears the advantage of adapting the threshold to an estimated distribution of reads in the sample. The last cut-off threshold is a fixed approach that removes the molecular features represented in a sample by less than 100 reads (hereafter “fixed 100 reads”), thus mimicking studies where exclusion thresholds are based on reads found in sequencing blanks (e.g. Macgregor et al., 2019).

**Table1:** List of published studies subjected to review, including details on referencing, type of used samples in the metabarcoding analysis, the organisms from which the pollen samples were collected, the type of cut-off threshold with a brief explanation of the threshold actually applied. Additional information are on the DNA barcode marker(s) utilized and on the utilization of the non-filtered sample composition tables in this study.

| Source | Type of sample | Organism | Cut-off threshold type | Detail on utilized cut-off threshold | DNA barcode marker(s) | Non-filtered dataset used in this study |
| --- | --- | --- | --- | --- | --- | --- |
| Baksay et al., 2020 | Mock pollen samples | - | Mixed | Sequences with a count of $\leq 10$, with no variants and with a count $< 5\%$ of their own count | ITS1, trnL | |
| Bansch et al., 2020 | Pollen from legs | *Apis mellifera*,<br>*Bombus spp.* (Apidae) | Not specified | Threshold on read count not reported | ITS2 |  |
| Bell et al., 2017 | Mock pollen samples | - | Negative controls | Removed identifications occurring at a lower frequency than identifications obtained in either negative control (isolation negative control 34 reads, PCR negative control 30 reads) | ITS2, rbcL |  |
| Bell et al., 2017(b) | Pollen from the whole body | Hymenoptera: Anthophila | Negative controls | Removed taxonomic classifications recorded from fewer reads than the maximum read number from a negative control (between 21-936 rbcL and 42-1124 ITS2) | ITS2, rbcL | X |
| Bell et al., 2019 | Mock pollen samples | - | Negative controls | Threshold based on the maximum sequence count from any negative control (11 and 34 ITS2, 8 and 30 rbcL) | ITS2, rbcL | X |
| Beltramo et al., 2021 | Honey | *Apis mellifera* (Apidae) | Proportional | OTUs with <0.2% of the reads | trnL |  |
| Biella et al., 2019 | Pollen from legs | *Bombus terrestris* (Apidae) | Variable: statistical-based | Receiver Operating Characteristics | ITS2 | X |
| Danner et al., 2017 | Pollen from legs | *Apis mellifera* (Apidae) | Proportional | Species <1% of the relative reads abundance per sample | ITS2 |  |
| deVere et al., 2017 | Honey | *Apis mellifera* (Apidae) | Not specified | Threshold on read count not reported | rbcl | X |
| Elliott et al., 2020 | Pollen from legs or scopa | Hymenoptera: Apidae, Halictidae, Megachilidae, Colletidae | Proportional | Taxa <1% of the proportion of all reads per plant taxon for each bee species | rbcl |  |
| Fahimee et al., 2021 | Pollen from the whole body | *Heterotrigona itama* (Apidae) | Fixed - Not proportional | OTUs with <2 reads | trnL |  |
| Galliot et al., 2017 | Pollen from the whole body | Diptera, Hymenoptera, Coleoptera, Lepidoptera | Negative controls | Based on control samples: 3 reads/genus per sample | ITS2 |  |
| Gous et al., 2018 | Pollen from the scopa | *Megachile venusta* (Megachilidae) | Proportional | Taxa <0.1% of total reads number per sample | ITS1, ITS2, rbcl |  |
| Gous et al., 2021 | Pollen from the scopa | *Megachile* spp. (Megachilidae) | Proportional | Taxa <0.1% of total reads number per sample | ITS2 |  |
| Hawkins et al. 2015 | Honey | *Apis mellifera* (Apidae) | Fixed - Not proportional | Taxa <10 sequences | rbcl |  |
| Jones et al., 2021 | Honey | *Apis mellifera* (Apidae) | Fixed - Not proportional | Singletons discarded | ITS2, rbcl | X |
| Khansaritoreh et al., 2020 | Honey | *Apis mellifera* (Apidae) | Not specified | Threshold on read count not reported | ITS2, rbcl |  |
| Leidenfrost et al., 2020 | Pollen from legs | *Bombus terrestris* (Apidae) | Not specified | Threshold on read count not reported | ITS2 |  |
| Lucas et al., 2018 | Pollen from the whole body | Syrphidae | Not specified | Threshold on read count not reported | rbcl |  |
| Lucas et al., 2018 (b) | Pollen from the whole body | Syrphidae | Not specified | Threshold on read count not reported | rbcl |  |
| Lucek et al., 2019 | Honey | *Apis mellifera* (Apidae) | Fixed - Not proportional | 5 reads per cluster (97% similarity) | ITS2 | X |
| MacGregor et al., 2019 | Pollen from proboscis | Lepidoptera (moths) | Negative controls | Read depth of 50 reads based on positive and negative controls | rbcl |  |
| Nünberger et al., 2019 | Pollen from legs | *Apis mellifera* (Apidae) | Not specified | Threshold on read count not reported | ITS2 |  |
| Peel et al., 2019 | Pollen from legs | *Apis mellifera*, *Bombus* spp. (Apidae) | Proportional | Species <1% of the total assigned long reads per sample | Genomic DNA |  |
| Piko et al., 2021 | Pollen from the whole body | *Bombus terrestris*, *B. pascuorum*, *B. lucorum* (Apidae) | Mixed | <100 reads per sample and removed species <1% of the sample's read count | ITS2 |  |
| Pornon et al., 2016 | Mock pollen samples, Pollen from whole body | *Hippaestrum* sp., *Chrysanthemum* sp., *Lilium* sp. ; Diptera, Hymenoptera, Coleoptera, Lepidoptera | Mixed | <1% of the most common sequences and <10 reads per sample | ITS1, trnL |  |
| Pornon et al., 2017 | Pollen from the whole body | Diptera, Hymenoptera, Coleoptera, Lepidoptera | Fixed - Not proportional | Sequences in number <1000 | ITS1, trnL |  |
| Pornon et al., 2019 | Pollen from the whole body | Syrphidae, Empididae, Apidae | Fixed - Not proportional | Sequences in number <1000 | ITS1, trnL |  |
| Potter et al., 2019 | Pollen from the whole body | Hymenoptera: Anthophila | Not specified | Threshold on read count not reported | rbcl |  |
| Richardson et al. 2015 | Pollen from legs | *Apis mellifera* (Apidae) | Not specified | Threshold on read count not reported | ITS2 |  |
| Richardson et al. 2015(b) | Pollen from legs | *Apis mellifera* (Apidae) | Fixed - Not proportional | Families found with only 1/3 amplicon libraries per sample | ITS2 |  |
| Richardson et al., 2019 | Pollen from legs | *Apis mellifera* (Apidae) | Proportional | Genera identified with only one marker and taxa with proportion of sequences <0.01% | ITS2, rbcL, trnL, trnH |  |
| Richardson et al., 2021 | Pollen from legs | *Apis mellifera* (Apidae) | Proportional | Genera identified with only one marker and with <0.001 proportional abundance of sequences | ITS2, rbcL, trnL |  |
| Sickel et al. 2015 | Pollen from nest | *Osmia bicornis*, *O. truncorum* (Megachilidae) | Proportional | Taxa <0.1% of reads per sample | ITS2 |  |
| Simanonok et al., 2020 | Pollen from legs | *Bombus affinis* (Apidae) | Mixed | <10 reads per OTU and taxa with <2% reads per sample | ITS2 |  |
| Smart et al., 2017 | Pollen from legs | *Apis mellifera* (Apidae) | Fixed - Not proportional | Taxa <50 reads | ITS1, ITS2 |  |
| Suchan et al., 2019 | Pollen from the whole body | *Vanessa cardui* (Lepidoptera) | Fixed - Not proportional | <100 reads per plant species per sample | ITS2 |  |
| Swenson & Gemeinholzer, 2021 | Mock pollen samples | - | Mixed | Taxa <0.1% of the sample reads of ITS1 and ITS2; removed identifications occurring at a lower frequency than identifications obtained in either negative control for rbcL | ITS1, ITS2, rbcL |  |
| Tanaka et al., 2020 | Pollen from honeycomb | *Apis mellifera* (Apidae) | Not specified | Threshold on read count not reported | rbcL |  |
| Tremblay et al., 2019 | Pollen from legs | *Apis mellifera* (Apidae) | Fixed - Not proportional | Taxa <100 reads | ITS2 |  |
| Vaudo et al., 2020 | Pollen from nest | *Osmia cornifrons* (Megachilidae) | Proportional | Taxa <1% relative read abundance and genera <0.3% of the total read counts per site across all sites | ITS2 | X |
| Wilson et al., 2021 | Pollen from nest | *Tetragonula carbonaria* (Apidae) | Proportional | Taxa identified in blank controls with abundance <1% of relative read abundance in real sample | ITS2, rbcL |  |

For each dataset, the variation in pollen species composition and species richness (standardized for the maximum number of species observed in a sample) for each sample was evaluated in response to the type of filtering used (i.e. no cut, proportional 1%, fixed 100 reads, and statistical ROC). Moreover, network indices describing the interactions between plants and pollinators were calculated through the R-package Bipartite and rnetcarto (Dormann, Gruber, & Fründ, 2008; Doulcier & Stouffer, 2015) for those datasets originated from studies based on insects direct characterization (specifically excluding one study on mock samples, Bell et al., 2019, and one study not clearly comparable with the other selected for network indices calculation because of the sample size and the experimental design, deVere et al., 2017). Specifically, the following community level indices were calculated: Connectance (i.e., measure of proportion of possible links actually recorded), Modularity (i.e., measure of the division of species into compartments, or modules, where species within modules share more interactions with each other than they do with species from other modules), and Shannon entropy (i.e., a measure of the diversity and complexity in the interactions of a species). Furthermore, at the level of a single individual pollinator, the connectivity index was calculated. This index quantifies the putative central role of an individual or of a species while connecting different parts of the whole network (Biella et al., 2017).

To evaluate changes in the pollen species composition of samples in response to the applied cut-off thresholds, we used distance matrices (jaccard distance) for an analysis of variance that uses permutations test with pseudo F-ratio (Andeson 2001) through the “adonis” function with R-package Vegan (Dixon 2003). Each dataset was analysed independently. The effect of the different cut-off thresholds on species richness was evaluated through a Generalized Linear Mixed Model (GLMM) approach with species richness as response variable and the type of filtering used (i.e. no cut, proportional 1%, fixed 100 reads, and statistical ROC) as covariate. The identity of the pollinator insect nested within the dataset was set as a random effect. Changes in interaction indices, both at the network and the individual level, were also evaluated through either a Linear Mixed Model or GLMM depending on the distribution and range of the response variable, with the type of filtering used (i.e. no cut, proportional 1%, fixed 100 reads, and statistical ROC) as covariate, and the dataset as random effect. The individual level connectivity was analysed as response variable, the type of filtering used (i.e. no cut, proportional 1%, fixed 100 reads, and statistical ROC) as covariate in interaction with the normalised degree of the pollinator individuals which was calculated as the number of plant species found in each sample divided by the overall number of plants in a given community. In this case, the sample identity nested within the dataset was included in the model as a random effect. For all the mentioned analyses, a comparison between the type of filtering used (i.e. no cut, proportional 1%, fixed 100 reads, and statistical ROC) were performed through a post-hoc test (Tukey’s HSD test). All the statistical analyses explained above were carried out with R (Version 3.6.1;R CoreTeam2019).

## 3 - RESULTS

### 3.1- TO PRUNE OR NOT TO A POLLEN DNA METABARCODING OUTPUT? A LITERATURE OVERVIEW

Overall, 42 research articles on pollen DNA metabarcoding were found and reviewed concerning the pruning of false positive or rare occurrences, and specifically the type of cut-off threshold applied (Proportional, Fixed not proportional, Variable: statistical based, based on Negative controls, Mixed, or Not specified). Furthermore, the analysed type of sample (i.e., honey, pollen mock samples, or pollen recovered from the body of insects, specifying the target portion of the body) and the organism from which the pollen was recovered were considered. Brief details on the applied cut-off threshold have also been considered, along with the employed DNA barcode marker and the availability of non-filtered sample composition tables subsequently involved in our analysis (Table 1). Concerning the strategies for managing false positives and rare occurrence, about one quarter of studies did not exclude false positives, while the remaining ones applied at least a pruning type. Specifically, the proportional cut-off thresholds was the most commonly applied, and involved 11 (26%) of the studies out of the panel of 42. Among these, the cut-off threshold calculated as 1% of the number of reads produced by each sample was the most recurrent. Only one study used a statistical approach (i.e., the ROC curve; Biella et al., 2019) to set a proportional cut-off threshold. Ten (24%) of the studies used a fixed number of reads chosen arbitrarily as cut-off threshold (e.g 100 or 1000 reads), and five (12%) used the number of reads produced by negative controls to set the threshold to remove false positives. Finally, five studies (12%) used a mixed approach that involved more than a single method to remove false positives. All of these details, along with a brief explanation of the strategies applied to set the false positive cut-off threshold for each of the reviewed studies are reported in Table 1.

These studies adopted DNA metabarcoding to address a range of cases. Among these, 27 studies (64%) recovered the pollen samples from the whole insect’s body or from specific body portions such as *scopa* and *corbiculae*. Four studies (10%) focused on the pollen stored in cavity nests or in hives, while five (12%) investigated mixed pollen mock samples to address methodological issues (optimization of DNA extraction or quantitative use of DNA metabarcoding reads). Finally, six studies (14%) analysed the taxonomic composition of honey, by looking at the pollen grains contained in it.

The vast majority of these studies (64%) relied on the ITS2 marker as a barcode region for species identification, although in some cases (26%), it was also combined with other barcode loci (e.g., rbcL).

### 3.2 EVALUATING THE CONSEQUENCES OF THE PRUNING TYPE

From the 42 reviewed studies, seven non filtered publicly available dataset were retrieved along with one unpublished dataset, produced by the authors, that was also included in the subsequent analysis (available upon request at http://doi/10.6084/m9.figshare.13637576, this data will be published after paper acceptance, ndr). Among these, four datasets were obtained by processing pollen found in nests or carried on insects bodies (Bell et al., 2017; Biella et al., 2019; Vaudo et al., 2020; along with the unpublished one, hereafter Tommasi et al., unpublished, that contains information about interactions between 249 insects and 156 plants). Three dataset comes from honey sample analysis (Lucek et al 2019; DeVere et al., 2017; Jones et al., 2021), and one was obtained from the analysis of pollen mock samples specially constructed for methodological assessments (Bell et al., 2019).

The comparisons among the effects of different pruning types on the community (pollen plant species) composition are summarized in Table 2, showing significant changes in the pollen species composition. Specifically, the main differences occurred between the no-cut and all the threshold-based prunings in all datasets (Table 2). Only minor community composition changes among threshold-based prunings were only occasionally found (Table 2)

**Table 2:** Comparison of the selected cut-thresholds on sample pollen species composition of several datasets based on analysis of variance based on permutations test with pseudo-F ratio. Dataset names (entitled with main author and year, see Table 1 for further details) are reported in the first column “Dataset”. The column “F” reports the pseudo-F ratio value and P the associated significance (α=0.05). Columns “A” to “F” report P value for the comparison between the different applied thresholds.

| Dataset | F | P full model | A<br>(Proportional 1% vs No cut) | B<br>(Fixed 100reads vs No cut) | C<br>(Proportional 1% vs Fixed 100reads) | D<br>(Statistical ROC vs no cut) | E<br>(Statistical ROC vs 1%) | F<br>(Statistical ROC vs Fixed 100 reads) |
| --- | --- | --- | --- | --- | --- | --- | --- | --- |
| Tommasi et al., (unpublished) | 0.819 | 0.806 | 1 | 1 | 1 | **0.031** | 0.314 | **0.045** |
| Bell et al., 2017 | 87.264 | **0.001** | **0.001** | **0.001** | 0.14 | **0.001** | **0.001** | **0.001** |
| Bell et al., 2018 | 39.817 | **0.001** | **0.001** | **0.001** | **0.01** | **0.001** | 0.658 | **0.001** |
| Biella et al., 2019 | 29.671 | **0.001** | **0.001** | **0.001** | 0.035 | **0.001** | 0.725 | **0.008** |
| Jones et al., 2021 | 6.538 | **0.001** | **0.001** | **0.001** | 0.944 | **0.001** | 0.233 | 0.855 |
| Lucek et al., 2019 | 5.465 | **0.001** | **0.001** | **0.001** | **0.038** | **0.001** | 0.975 | **0.058** |
| deVere et al., 2017 | 2.415 | **0.024** | **0.004** | 0.212 | 0.538 | **0.003** | 0.704 | 0.152 |
| Vaudo et al., 2020 | 11.553 | **0.001** | **0.001** | **0.001** | 0.556 | **0.001** | 1 | 0.578 |

Plant species richness of pollen from samples was found to be significantly influenced by the pruning type (χ_3_^2^= 468.22, p < 0.001). Specifically, consistently higher species richness per sample was found in the unfiltered (no cut), compared to all the other pruning types (proportional 1%, fixed 100 reads, and statistical ROC). A significant difference between the proportional 1% and the statistical ROC approaches was also found, with the latter reducing species richness more (Fig1a, Table 3).

**Fig 1:**
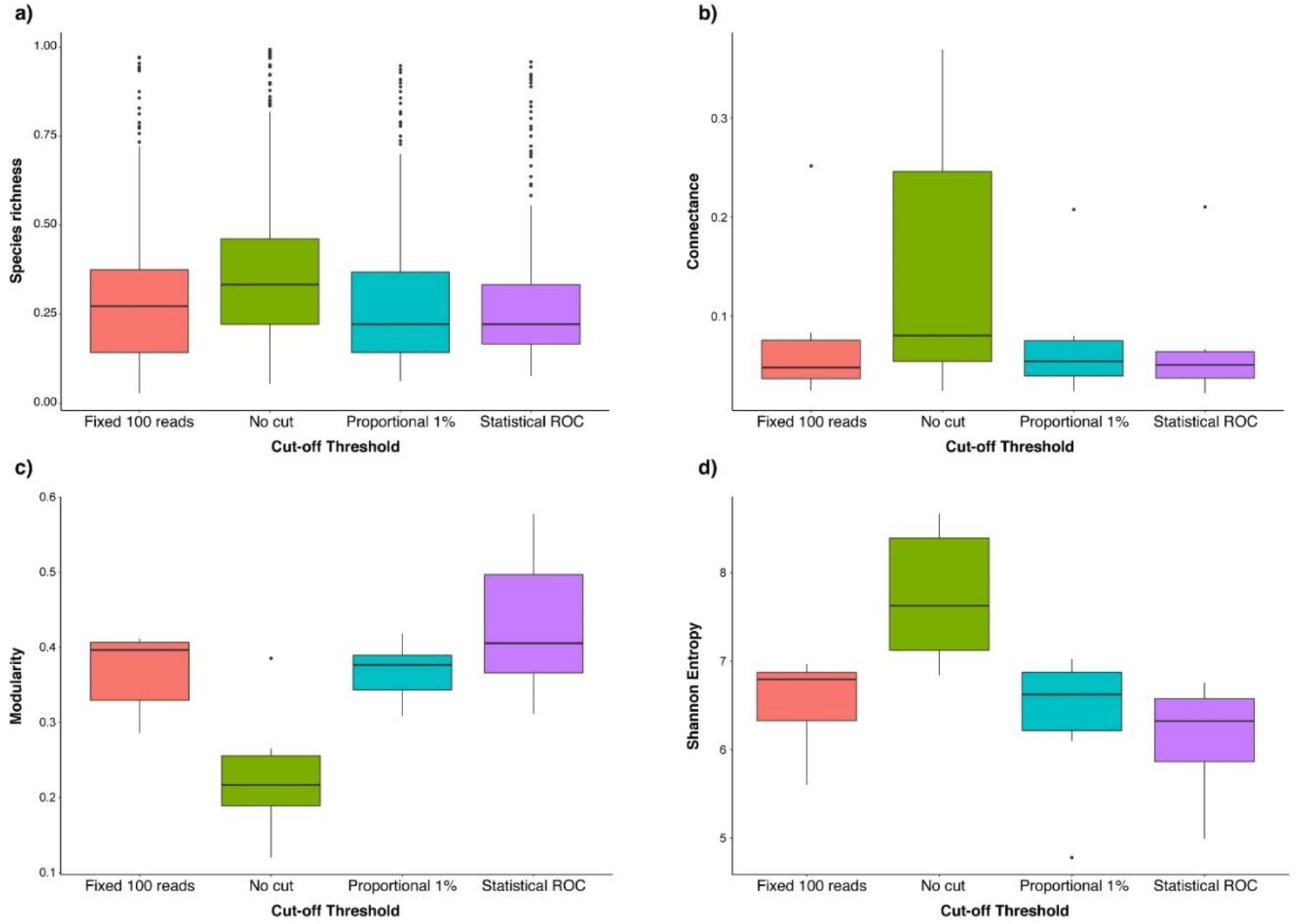
Variation of Species richness (a), Connectance (b), Modularity (c), and Shannon entropy (d) using different cut-off thresholds (i.e No cut, Fixed 100 reads, Proportional 1%, Statistical ROC)

**Table 3:**
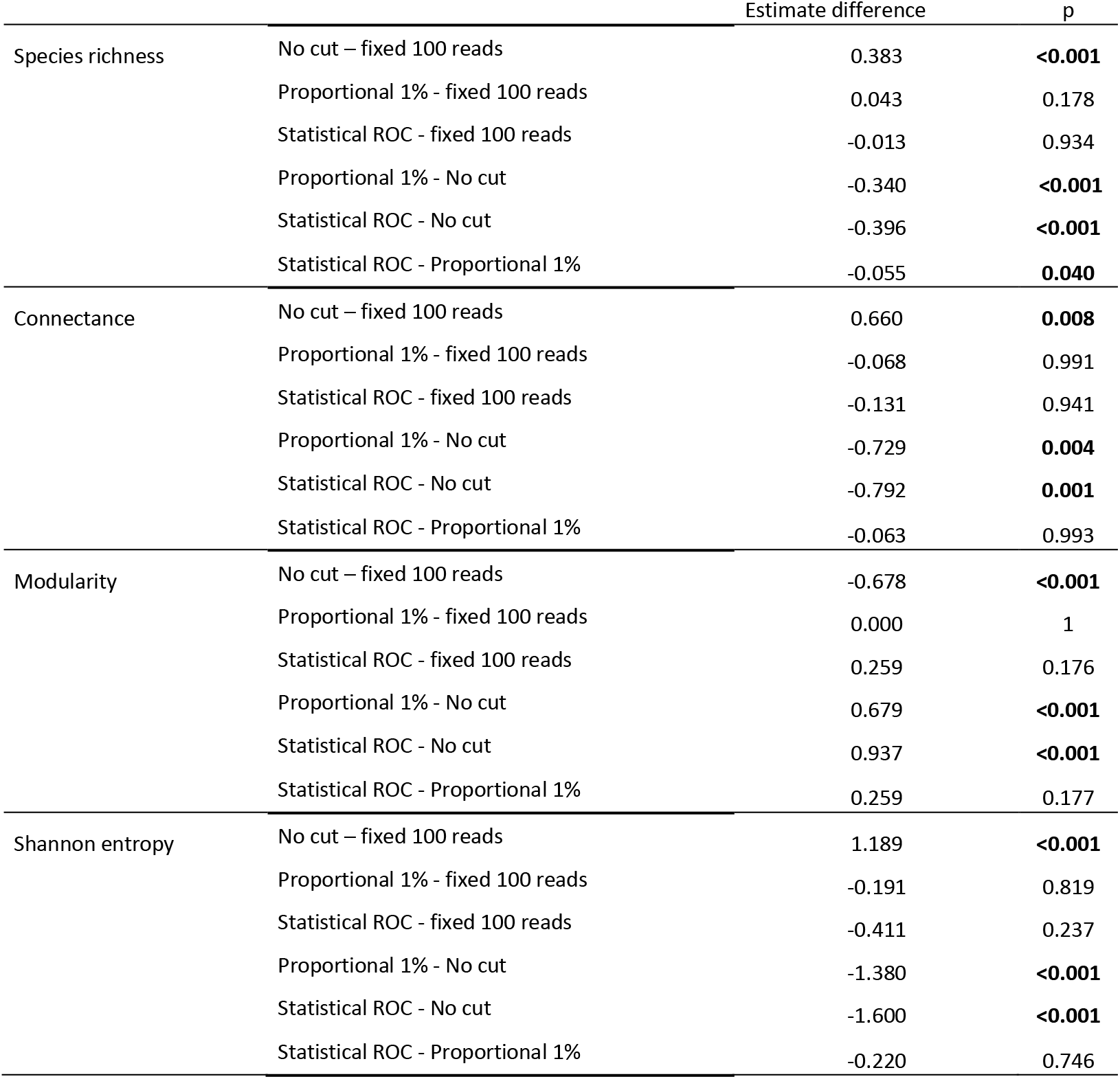
Statistical comparison of the selected cut-off thresholds (i.e. No cut, Fixed 100 reads, Proportional 1%, Statistical ROC) on Species richness, Connectance, Modularity, and Shannon entropy (Tukey’s multiple comparison test, α = 0.05).

The effects of pruning type on the network level indices were significantly found on connectance (χ_3_^2^ = 11.642, p = 0.008), modularity (χ_3_^2^ =25.273, p < 0.001) and Shannon entropy (χ_3_^2^ = 29.907, p < 0.001). Connectance (Fig1b, Table 3) and Shannon entropy indices (Fig1d, Table 3) were significantly higher while Modularity significantly lower (Fig1c, Table 3) in the unfiltered (no cut) compared to all the other pruning types (proportional 1%, fixed 100 reads, and statistical ROC). In addition, in most cases, these indices after ROC pruning were notably different from values obtained from “proportional 1%” and “fixed 100 reads” networks (Fig 1c; Fig 1d).

The individual level index of connectivity showed a significant effect of the interaction between the applied pruning and the normalised degree index (χ_3_^2^ = 609.2, p < 0.001). Specifically, the connectivity was lower in the unfiltered (no cut) compared to all the other pruning types (proportional 1%, fixed 100 reads, and statistical ROC) for any value of the normalised degree (i.e., both for generalist and for specialist individual pollinators, Fig 2, Table 4).

**Fig 2:**
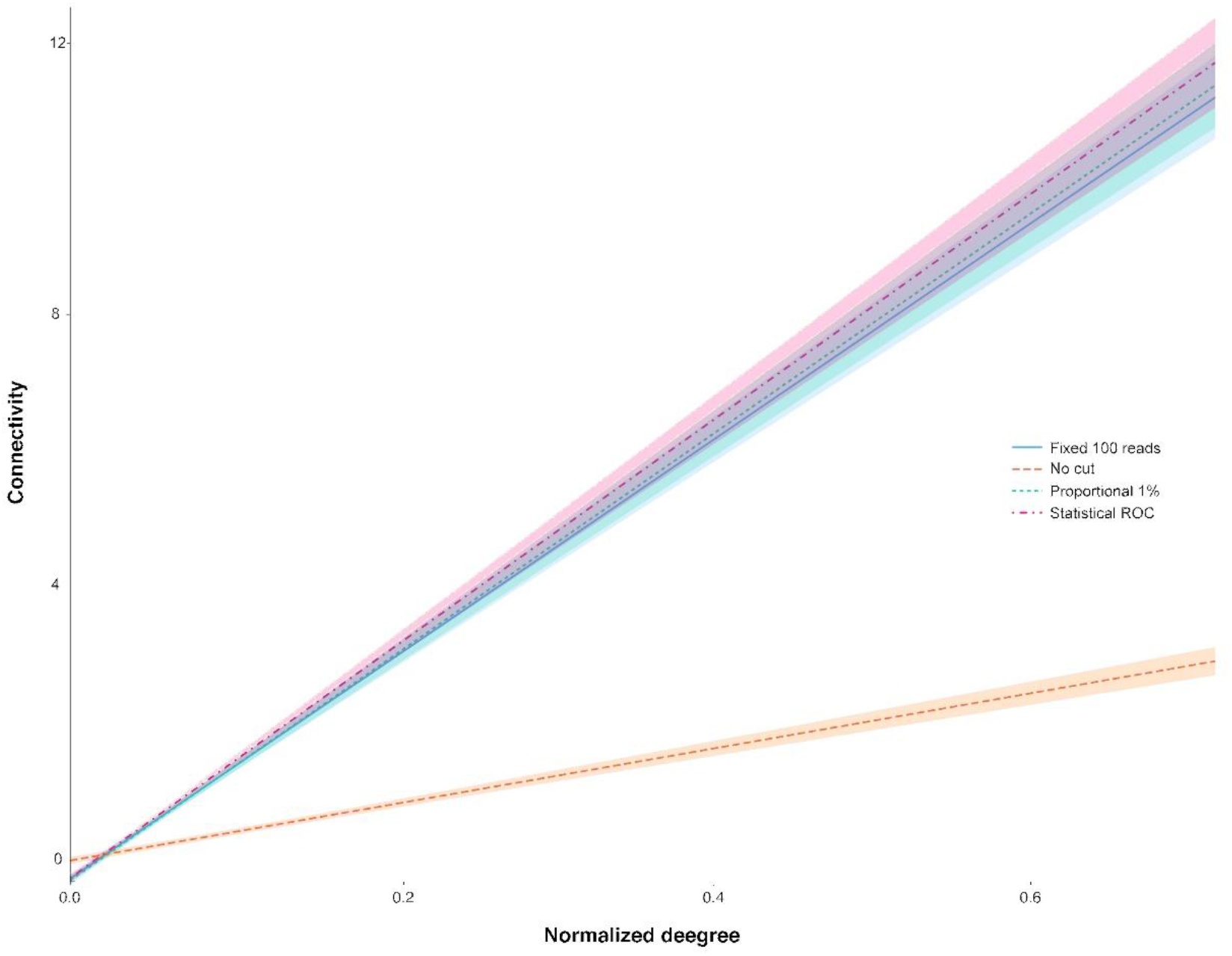
Response of the species level index connectivity to the cut-off threshold-based treatments (i.e No cut, Fixed 100 reads, proportional 1%, statistical-based ROC) in interaction with the individuals normalised degree.

**Table 4:** Statistical output related to the comparison of the selected cut-off thresholds in interaction with individuals’ normalized degree on the individual pollinator index Connectivity (Tukey’s multiple comparison test, α = 0.05).

| Cut-off threshold: normalised degree<br>comparison | Estimate difference | p |
| --- | --- | --- |
| No cut – fixed 100 reads | -11.896 | **<0.001** |
| Proportional 1% - fixed 100 reads | 0.279 | 0.756 |
| Statistical ROC - fixed 100 reads | 0.694 | 0.456 |
| Proportional 1% - No cut | 12.175 | **<0.001** |
| Statistical ROC - No cut | 12.590 | **<0.001** |
| Statistical ROC - Proportional 1% | -0.414 | 0.662 |

## DISCUSSION

Since its ‘formalization’ in 2012, the DNA metabarcoding approach has revolutionized the field of biodiversity investigation and provided many insights into the study of biological interactions. Its application has rapidly spread and has contributed to support several research contexts such as microbiome (Bruno et al., 2018; Frigerio et al., 2020), food, (Galimberti et al., 2015; 2019; 2021), trophic ecology (Casey et al., 2019; Arrizabalaga-Escudero et al., 2018), and environmental DNA-based analyses (Ruppert et al., 2019). In spite of its usefulness, the whole DNA metabarcoding pipeline and specifically the bioinformatic processing could deeply influence the obtained results and their interpretation (Andriollo et al., 2019; Zinger et al., 2019 Elbrecht et al., 2017). Therefore, in this study we attempted to evaluate the effects of the adopted pruning approach used to filter the reads obtained through HTS. Specifically, we focused on the analysis of pollen DNA metabarcoding data in the framework of plant-insect interactions, being aware that the outputs of our investigation could be extended to the other typologies of DNA metabarcoding-based studies. Although the issue of removing false positives and rare species is quite neglected in the literature regarding the adopted bioinformatic DNA metabarcoding pipeline (but see Ficetola et al., 2016), the choices made when analysing a HTS output could generate relevant effects on the obtained community composition, species richness and species interactions. These aspects would deeply impact the ecological outcomes of the investigated experimental system.

Reliable DNA metabarcoding outputs from pollen analysis require robust and replicable approaches for treating HTS molecular features (e.g.,ESVs and OTUs) that should be coherent and comparable among different studies. However, our literature overview highlighted a high heterogeneity in the type of pruning adopted to remove false positives and infrequent species. This is particularly appreciable even among studies that focused on similar analytical matrices (e.g. pollen from insect’s body, pollen from cavity nests and honey).

Our literature review shows that the proportional approach emerges as the most recurrent, that is to remove those molecular features/species under a certain proportion of the total reads per sample. This is not surprising, as it is an approach also well represented in other DNA metabarcoding-based studies (e.g., Bohmann et al., 2018; Arribas et al., 2021 and Casey et al., 2019). This could be explained by the ease of calculating proportions, and by the advantages of using these rather than fixed thresholds (e.g., comparison among different samples and a low filtering impact in the case of samples with low number of total reads). However, in the case of pollen DNA metabarcoding data, we found no concordance between different authors about the exact amount of proportion of reads to be used as threshold, and the reason under the choice of a particular percentage (e.g. Sickel et al., 2015 used 0.1% while Richardson et al., 2019 used 0.01%; see Table 1). However, it should be noticed that Peel et al. (2019), while analysing mock pollen samples with known composition, highlighted that false positive identified in samples occurred at a rate lower than 1%, thus supporting this filtering strategy. On the other hand, caution should be recommended prior to generalising the 1% threshold as a universally effective filtering practice, as for samples with extremely high total reads it might be better to use a lower value.

The second most recurrent cut-off approach is based on a fixed number of read counts, used as a uniform threshold among all samples (e.g. 50 as in Smart et al., 2017, 100 as in Tremblay et al., 2019 see Table 1), the most frequent amount being 100 reads per sample. As reported above, also with this approach the specific amount of reads chosen as the cut-off threshold is usually poorly supported by clear biological reasons. Studies using higher threshold values would remove false positives and truly occurring taxa in excess. An example of this is Pornon et al. (2017) and (2019), who observed how a threshold of 1000 reads per plant species ensures the removal of the vast majority of grass pollen species. Despite speculating that the source of that grass pollen was airborne contamination, the species that pollen belonged to, actually occurred at the study areas and thus those species shall be considered true positives. Therefore, lower or higher cutting values might be chosen based not only on removing false positives, but also potential environmental contamination or infrequent species. Conversely, in other studies, the choice of the cut-off value is clearer and derives from the use of sequenced negative controls. In this case, the maximum number of reads found in blank samples is set as the threshold for the false positive removal (see Table 1). The rationale behind this approach is that it should allow removing false positives exclusively originating from laboratory activities (Bell et al., 2019). It is not clear how effective this method is when it is adopted to remove rare, infrequent, ecologically unmeaningful species obtained from HTS processing.

Unexpectedly, the literature survey (Table 1) showed that nearly a quarter of pollen DNA metabarcoding studies did not filter raw data and thus did not remove possible false positives from the analysed samples. This methodological choice can hardly be supported by biological reasons or particular research constraints. Removing false-positives is a priority, and the analyses we performed here clearly suggest that filtering the HTS output with a cut-off threshold leads to significant differences compared to the unfiltered output matrix, especially in species composition, species richness and plant-pollinator interactions. This indicates that a “no cut” strategy could deeply impact the ecological interpretation of results and could lead to misleading conclusions.

Unfiltering could deceive results, and specifically, the application of any of the cut-off thresholds here investigated shape the community composition and also decrease the species richness in comparison to non-filtered data. These results are even amplified by the research aims. In studies related to the characterization of insects foraging behaviours, for example, un-filtering could overestimate the number of plants foraged by a pollinator, and obviously to the wrong assessment of generalism, foraging niche, and of delivered pollination ecosystem service. In studies on honey composition, a no cut strategy could mislead about the purity of products, with consequences that could involve commercial issues.

In our simulations, the reads filtering impacted not only species composition and richness, but also the ecological networks associated with each non-filtering and filtering strategy. The significant variation of network indices in response to the applied cut-off thresholds, both at the community and the individual levels, further confirms how the ecological outputs of DNA metabarcoding studies are influenced by the strategy of rare species and false positive removal. Specifically, greater differences occurred when comparing index values calculated from filtered (Fixed 100 reads, proportional 1%, and statistical ROC) and non filtered (no cut) data. The implications of the changes in network indices are very high, as for instance network entropy, connectance, modularity and connectivity refer to the network stability and resilience, to the ability to buffer perturbations and to the stabilizing role of central hub species (Tylianakis et al., 2010; Thébault & Fontaine, 2010; Biella et al., 2017; Strydom et al., 2020), since the higher the index difference between filtering strategy, the higher the potential for misleading ecological results. In particular, filtering decreased the network’s connectance and entropy indexes. This result aligns well with the recorded lower species richness per sample in filtered dataset and it can be explained by an overall decrease in network number of realized links (i.e., less plant species found on pollinator bodies or samples). In other words, by decreasing the numerosity of links, filtering likely yields networks with slightly higher element-specific linkage compared to networks deriving from a non filtering approach. Moreover, filtering increased modularity and connectivity of networks. This result further clarifies that filtering decreases the ubiquity of links among elements (less features connecting to everyone), thus allowing for better emergence of ordered patterns of well-defined compartments of interactions and important hub species connecting them. Hence, in practice, unfiltering returns networks richer in links, which are even more ubiquitous among elements, with the high potential of overestimating foraging strategies and network resilience. Among the filtering strategies analysed here, the statistical ROC approach appears to be the most conservative one, since it tends to yield the lowest species richness, the highest modularity, and the lowest connectance and entropy. Thus, it might remove not only the false positives from samples, but also the infrequent and thus unmeaningful species. It should be noticed that ecological patterns emerging or confirmed even in a conservative framework are more likely to be trustworthy. Even if this approach has rarely been applied in the HTS pollen literature to date (Table 1), it was specifically developed to distinguish “true signals” from “noise” (Fan et al., 2006) in molecular biology-based studies and could constitute a promising avenue for the interpretation of pollen DNA metabarcoding data (Biella et al 2019). For instance, it has been used in other DNA-based research fields (Nutz et al., 2011, Siddique et al., 2021), such as for eDNA where it is proved to increase the reliability of data (Serrao et al., 2018). Because it is a conservative approach, ROC may be favoured in studies willing to highlight ecologically meaningful species composition, richness and interactions, while sacrificing the pursuit of high species richness based on keeping elements of rarity, potential contaminants and false positives.

In conclusion, our survey should help in improving the awareness on the use of DNA metabarcoding for pollen identification and its applications in a broad spectrum of ecological and biological research. To date this powerful tool still requires the development of shared approaches to provide reliable, repeatable, and comparable data, since high heterogeneity emerges from the scientific literature on the bioinformatics filtering of molecular features obtained from HTS outputs. In particular, we recommend that researchers may (i) always make both raw unfiltered and filtered data easily accessible, thus improving the possibility of exploring large amounts of data, and consequently the growing rate of human knowledge in strategic research fields such as pollination ecology. The authors may also (ii) apply (and Journals’ reviewers may encourage for) filtering from false-positives and possibly also from infrequent species although depending on research aims. Moreover, (iii) the specific type of filtering shall be clearly justified under a biological perspective, also evaluating the efficiency and universality of the loci selected for species identification and the consequent taxonomic resolution of molecular feature assignments. Moreover, (iv) the specific strategy might be decided based on whether the research aim would benefit from or need a conservative approach. This is the case, for example, of using DNA metabarcoding reads as presence/absence data, an approach that leads to large difference in species detection in case false positives and rare species are non removed from the assignments (Ficetola et al., 2015; Deagle et al., 2018); if so, the ROC filtering should be preferably adopted. Otherwise, it is always recommended to apply a filter either based on a percentage with a clear biological support or rather based on a fixed value like sequencing blanks, thus excluding only false positives while keeping environmental contamination and infrequent species.

By analysing available dataset from pollen DNA metabarcoding we proved that all the possible false positive removal strategies affect sample composition and consequently the ecological interpretation that could be extracted by them. This emphasizes the need to firstly adopt and secondly clearly justify the choice behind the adoption of different criteria for false positives and rare occurrences removal, taking into account the research aims and the expected ecological outputs as well.

